# Enhancing anti-gastrointestinal cancer activities of CLDN18.2 CAR-T armored with novel synthetic NKG2D receptors Containing DAP10 and DAP12 signaling domains

**DOI:** 10.1101/2023.05.17.541124

**Authors:** Minmin Sun, Hongye Wang, Ruidong Hao, Youtao Wang, Yantao Li, Yunpeng Zhong, Shuangshuang Zhang, Bo Zhai, Yuanguo Cheng

**Affiliations:** School of Pharmacy, Fudan University, Shanghai, P.R. China; China State Institute of Pharmaceutical Industry, Shanghai, P.R. China; Suzhou Immunofoco Biotechnology Co., Ltd., Jiangsu, P.R. China; Department of Interventional Oncology, Renji Hospital, Shanghai Jiao Tong University School of Medicine, Shanghai, China; State Key Laboratory of Oncogenes and Related Genes, Shanghai Cancer Institute, Renji Hospital, Shanghai Jiao Tong University School of Medicine, Shanghai, China

**Keywords:** CAR-T, Solid tumor, CLDN18.2, NKG2D, Antigen Heterogeneity

## Abstract

Chimeric antigen receptor (CAR) T therapies have shown remarkable efficacy in hematopoietic malignancies, but their therapeutic benefits in solid tumors have been limited due to heterogeneities in both antigen types and their expression levels on tumor cells. NK group 2 member D ligands (NKG2DLs) are extensively expressed on various tumors and absent on normal tissues, making them a promising target for cellular immunotherapy. DAP10 and DAP12 function as adaptor proteins in NK cells to transduce activating signals, and recent studies have revealed DAP10 and DAP12’s additional role as a co-stimulatory signal in T cells. Our pre-clinical data showed that CAR-T targeting CLDN18.2 is highly effective in gastrointestinal (GI) cancers, but the heterogeneous expression of CLDN18.2 poses a treatment challenge. To complement this antigen deficiency, we demonstrated that NKG2DLs were extensively expressed in GI tumor tissues and formed an ideal dual target. Here, we reported a CLDN18.2 CAR design armored with synthetic NKG2D receptors (SNR) containing DAP10 and DAP12 signaling domains. This novel CAR-T showed improved cytotoxicity against tumor cells with heterogeneous expression of CLDN18.2. The possible underlined mechanism is that SNR promotes CAR-T memory formation and reduces their exhaustion, while also enhancing their expansion and ability to infiltrate immune-excluded tumors in vivo. Taken together, SNR with DAP10/12 signaling and their synergistic involvement, increased CAR-T function and overcame the antigen deficiency, providing a novel treatment modality for solid GI tumor.

## Introduction

Chimeric antigen receptors (CARs) are synthetic receptors that redirect immune cells to recognize tumor cells expressing targeted antigens in an MHC independent manner. CAR-T cell therapies targeting CD19 or BCMA have shown unprecedented efficacy in treating hematological malignancies^1–3^. However, the use of CAR-T cell therapies in solid tumors has been limited due to the absence of suitable targets that are highly and homogeneously expressed on tumor cells^4–7^ and the tumor environment suppression and CAR-T exhaustion and proliferation. This has resulted in a weak antitumor efficacy in solid tumors.

Claudin18.2 (CLDN18.2) has emerged as a promising solid tumor target. it is a stomach-specific isoform of CLDN18 which belongs to a tight junction protein family^8,9^. CLDN18.2 is highly expressed in cancer cells, particularly in gastric cancer/gastroesophageal junction (GC/GEJ) and pancreatic cancers, while minimally expressed in normal tissues except stomach^9–12^. Therapies targeting CLDN18.2 with antibody or CAR-T have demonstrated primary efficacy and good safety profiles in clinical trials^13–15^. Recently, CLDN18.2 targeted CAR-T therapy was evaluated in clinical trial CT041. For patients with GC, the ORR was 57.1% while it increased to 63% in patients with more than 70% CLDN18.2 expression^16^. Despite the high response rate, most patients had disease progression within 6 months including those with PR. One possible reason for this rapid progression was the outgrowth of target negative tumor cells, which might be due to the heterogeneity of CLDN18.2 expression in tumors. Tumor heterogeneity consists of intra-tumoral and inter-tumoral heterogeneity, intra-tumoral heterogeneity implies the inherent temporal-spatial differences between distinctive subpopulations of tumor cells, which heterogeneously express different markers^17^. Thus, designing CAR-T cells to target multiple antigens can overcome this heterogeneity so that the escape could be prevent.

One potential approach to overcome tumor heterogeneity and enhance the anti-tumor activity of CAR-T cells is to utilize the interaction between NKG2D/KLRP1, an activating receptor in natural killer (NK) cells, and its stress-induced ligands (NKG2DL), which include MICA, MICB, and ULBP1-6^18,19^. Under physiological conditions, NKG2D ligands were usually overexpressed on viral infected or DNA damaged cells but not expressed in healthy tissues^20,21^. However, human cancers including gastric adenocarcinoma can upregulate NKG2D ligand expression ^22–24^. DAP10 and DAP12 function as adaptor proteins and transport co-stimulatory signaling in both NK and T cells. A previous study suggested that expression of NKG2D in CD8^+^ T cells could favor the differentiation into central memory T cells and stem like memory T cells via DAP10 and DAP12 signaling in T cells^25^. Therefore, the harness of this interaction is an ideal approach to enhance the anti-tumor activity for CAR-T cell therapy.

In this study, we investigated the expression of CLDN18.2 and NKG2D ligands in human cancer micro-tissue array and showed that NKG2D ligands and CLDN18.2 were complementarily expressed in human gastric cancer, which favors NKG2D as an ideal target for dual targeting. Also, we proposed a novel design of CLDN18.2 CAR armored with synthetic NKG2D receptors (SNR) containing DAP10 and DAP12 signaling domains. Functionally, the SNR CAR-T have a higher memory T cell portions and showed longer persistence compared to CLDN18.2 CAR-T. Both in vitro and in vivo results showed that SNR CAR-T could eradicate not only CLDN18.2 or NKG2DL single positive tumors but could also repress growth of heterogeneous tumors. The novel structure indicates that SNR with DAP10/12 signaling and their synergistic involvement, increased CAR-T function and overcame the antigen deficiency, providing a novel treatment modality in resolving tumor heterogeneity.

## Methods and materials

### Isolation of CD3-positive T cells and construction of CAR-T

In this experiment, CD3 magnetic beads (Miltenyi Biotech) were used to sort CD3-positive T cells from peripheral blood mononuclear cells (PBMCs) and T cells were cultured in X-vivo (Lonza) medium supplemented with 5% FBS (Gibco), 100 mg/mL penicillin, 100 mg/mL streptomycin sulfate (Gibco), and 300 U/mL IL2 (Peprotech). After sorting with magnetic beads, CD3^+^ T Cells were stimulated with 10μg/ml anti-CD3 antibody and anti-CD28 antibody (Novoprotein) in six-well plates. After 24 hours, the activated T cells were infected with lentivirus containing CAR construct at a multiplicity of infection (MOI) of 10, and transduction rates were measured by flow cytometry at 72-hour post-activation.

### Cell lines and culture

The Human skin cancer cell line A431 and gastric cancer tumor cell line NUGC4-luc were provided by SHANG HAI MODEL ORGANISMS Co., Ltd. A431-CLDN18.2 cell line was generated by lentiviral infection of A431. All tumor cells were cultured with DMEM medium (Life Technologies) supplemented with 10% FBS, 100 mg/mL penicillin, and 100 mg/mL streptomycin sulfate in 37℃ humidified incubators with 5% CO2. All cell lines used in this study were authenticated using Short Tanderm Repeats (STR) analysis by the Shanghai Biowing Applied Biotechnology (Shanghai, China).

### In vitro cytotoxicity and cytokine secretion assays

We measured CAR-T cytotoxicity by detecting annexin-v positive tumor cells after co-culturing with CAR-T cells with FACS. Before co-culturing, different tumor cell lines were stained with carboxyfluorescein succinimidyl ester (CFSE) following the manufacture’s protocols and cultured in X-vivo (Lonza) supplemented with 5% FBS (Gbico) and 1% penicillin and streptomycin (Thermo) solution. 10,000 tumor cell lines were seeded into 96-well plate and then CAR-T were added with different effector: target ratio (3:1, 1:1, 3:1). After co-culturing for 5 hours, total cells were collected and cultured with APC-Annexin-V proteins for 20 min, finally, the mixed samples were analyzed by FACS. Meanwhile, supernatants from cell cultures were harvested for detecting cytokine using LEGENDplexTM Human Th1 Panel (5-Plex) (BioLegend). Samples were diluted 5-fold using assay buffer and then mixed with beads and shaken for 2 hours at 500 rpm in a 96 plate well. After washing with 1X washing buffer for twice, detection antibodies and streptavidin-phycoerythrin were added, and the plate was shaken for 1 hour and 30 minutes, respectively. Finally, beads were suspended with 200 μL PBS and mean fluorescence intensity (MFI) were detected with FACS.

### In vivo xenograft model

In this experiment, three kinds of cells were used to construct the xenograft model, including NUGC-luc, A431 and A431-18.2. Briefly, NSG mice were anesthetized with 3-4% isoflurane prior to inoculation. About 5 × 10^6^ cells were resuspended in PBS, mixed with an equal volume of Matrigel, and then inoculated into mice by subcutaneous injection in a volume of 200 µL. When the tumor grows to an average of about 100-150 mm^3^, mice were randomly divided into several groups and each group contains 6-8 mice. 2-3 mice in each group were euthanized and their tumor tissues were extracted, followed by fixation, and embedding. All animals were housed in a specific pathogen-free environment (12 h light/12 h dark with lights on at 7.00 h 21±2℃) with food and water ad libitum. This study was performed in strict accordance with institutional guidelines and approved by the institutional Animal Care and Use Committee of Shanghai Model Organisms. Here, immunofluorescence was used to analyze the infiltration of Car-T in tumor tissues.

### RNA sequencing

Total RNA was isolated from each CAR-T sample using the RNA minikit (Qiagen, Germany). RNA quality was examined by gel electrophoresis and with Qubit (Thermo, Waltham, MA, USA). For RNA sequencing, RNA samples from seven to nine biological replicates at each time point (0,12, 36 and 72h) were separated to three independent pools, each comprised of two or three distinct samples, at equal amounts. Strand-specific libraries were constructed using the TruSeq RNA sample Preparation kit (Illumina, SanDiego, CA, USA),and sequencing was carried out Using the Illumina Novaseq 6000 instrument by the commercial service of GenergIo technology Co.Ltd (Shanghai, China).The raw data was handled by Skewer and Data quality was checked by Fast QCv0.11.2 (http://www.bioinformatics.babraham.ac.uk/projects/fastqc/). The read length was 2×150bp.Clean reads were aligned to the Human genome hg38 using STAR. StringTie.The expression of the transcript was calculated by FPKM (Fragments PerKilobase of exon model perMillion mapped reads) using Perl. Differentially Expression transcripts (DETs) were determined using the MA-plot-based method with Random Sampling (MARS) modeling the DEGseq package between different time Points (12hptvs.0hpt,36hptvs.0hpt,72hptvs.0hpt). Generally, inMARS model, M=log2C1-log2C2, and A=(log2C1+log2C2)/2(C1and C2 denote the Count so reads mapped to a specific gene obtained from two samples). The Thresholds for determining DETsare P<0.05 and absolute fold change≥2. Then DETs were chosen for function and signaling pathway enrichment analysis using GO And KEGG database. The significantly enriched pathways were determined when P<0.05 and at least two affiliated genes were included.

### Flow cytometry

Flow cytometry was conducted following routine protocols. About 2×10^5^ cells were harvested, washed twice with PBS, then the antibody was mixed with the cell suspension at a ratio of 1:500 and incubated at room temperature for 20 minutes. All samples were then analyzed on a flow cytometer. In this study, the transduction rate of lentivirus on Car-T cells was analyzed by FITC-anti-VHH antibody (GenScript Inc.). Target cells were detected for NKG2D ligands using anti-MICA/MICB and anti-ULBP2/5/6. Phenotypes in Car-T cells were detected using anti-CD25, anti-CD69, anti-62L, anti-CD45RA, anti-PD-1, anti-CD27 and all antibodies are used for flow cytometry were purchased from Biolegend.

### Immunohistochemical (IHC) assay

Here, the samples for our IHC analysis are microarray purchased from Bioaitech Co., Ltd. The D046St01 microarray Contains 40 cases of gastric adenocarcinoma and 6 cases of adjacent gastric tissue. The main process of IHC analysis is sectioning, dewaxing, blocking, and staining. Sections were incubated with primary antibody at 4℃ overnight at a dilution ratio of 1:500, then sections were stained by horseradish peroxidase (HRP)-conjugated for 30 minutes at 37°C; The primary antibodies are anti-CLDN18.2 (Abcam), anti-MICA/MICB (Abcam), anti-ULBP1 (R&D), anti-ULBP2/5/6 (R&D) and anti-ULBP3 (R&D).

### Mass Cytometry

Antibody panel setup. Anti-VHH antibodies used to detect CAR-T cells were customized by Polaris Biology, China. The rest of the mass cytometry antibodies (CytoATLAS, Polaris Biology, China) are listed in supplementary Table 1.

Sample staining and acquisition. Cells were washed with LunaStain cell staining buffer (Polaris Biology, China) and first stained with Fc block (Biolegend, USA) for 10 min at room temperature. Cells were then stained with 10μL of Cisplatin reagent (Polaris Biology, China) and the heavy metal-labeled membrane antibody mixtures for 30 min at room temperature. Cells were washed twice and fixed in LunaFix cell fix buffer (Polaris Biology, China) for 5 min. Cells were then washed and resuspended in LunaPerm cell perm buffer (Polaris Biology, China) for 30 min. Cells were then washed and incubated with heavy metal-labeled intracellular antibodies mix for 1 h at room temperature. Cells then washed twice with cell perm buffer and stained with Ir-DNA intercalator reagent (Polaris Biology, China) for 10 min. After staining, cells were washed and adjusted to 1 million cells per milliliter in LunaAcq cell acquisition solution (Polaris Biology, China) together with 20μL of SureBits element calibration beads (Polaris Biology, China). Cell acquisition was performed at 300 events/ second on a mass cytometer (StarionX1, Polaris Biology, China).

Data analysis. After acquisition, mass cytometry data were normalized and converted into standard FSC 3.0 files (StarionX1, Polaris Biology, China). Manual gating was performed using FlowJo (BD Biosciences, USA). Uniform Manifold Approximation and Projection (UMAP) was used to get an overview of the immune compartment. To identify different cell subtypes, FlowSOM clustering and metaclustering was performed.

### Statistical analysis

All statistical tests were conducted with GraphPad (v8.0) and R software (v4.2.1). GraphPad Prism 8 was used for unpaired Student’s t test and two-way ANOVA test. Boxplots were represented as median and interquartile range, while bar plots were presented as means ± SEM. *P< 0.05 was regarded essential.

## Results

### Heterogeneous expression of CLDN18.2 limits anti-tumor efficacy of conventional single-targeting CAR-T

Claudin18.2 (CLDN18.2), a gastric-specific isoform of the tight junction protein of CLDN18, has been regarded as a potential therapeutic target for gastric cancer^8,9^. We evaluated the expression profile of CLDN18.2 in a human gastric cancer tissue microarray through immunohistochemistry. Consistent with previously report^9^, we found that CLDN18.2 was stained positive in only about 38% gastric cancer tissues. Among these positive cancer tissues, CLDN18.2 intensity showed a heterogeneous pattern of expression with some regions of low or negative staining (Fig.1A-C). To test whether NKG2DLs was an ideal dual target with CLDN18.2 in gastric cancer, we investigated the expression profile of NKG2DLs including ULBP1, ULBP2/3/5/6 and MICA/B in the same tumor tissues. As expect, we found that ULBP1 was highly expressed in most gastric tissues, whereas MICA/B, ULBP2/5/6 and ULBP3 were expressed by some tissues (Fig.1A). Further analysis showed that at least one NKG2D ligand was expressed in most gastric cancer tissues, and a total of 89% gastric cancers expressing some NKG2D ligands (Fig.1B), and that NKG2D ligands was positively expressed in CLDN18.2 negative tissues (Fig.1C). Thus, either CLDN18.2 or NKG2DLs was expressed in most gastric cancers (Fig.1C), which establishes our rationale to target both by the CAR-T.

**Fig.1.**
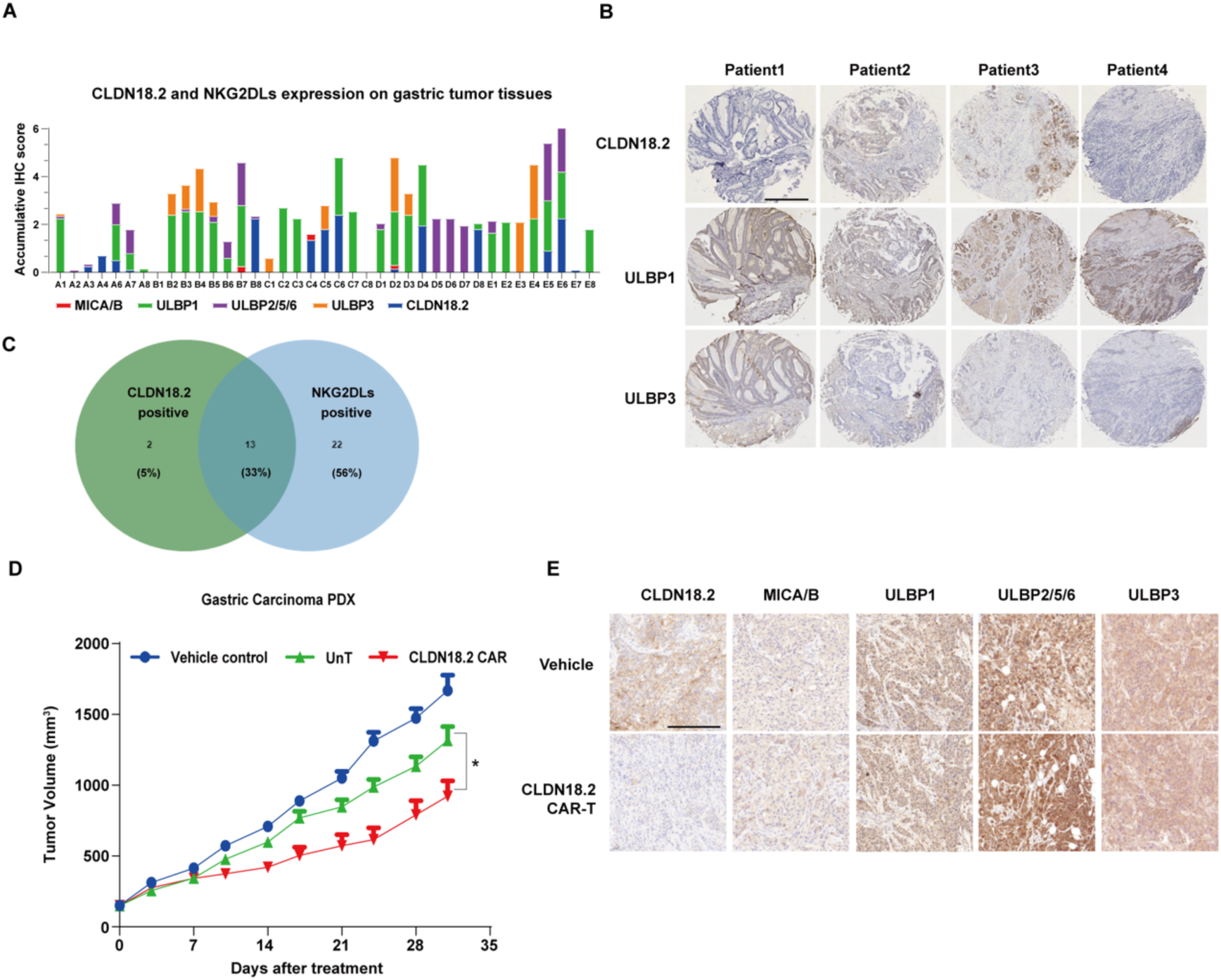
CLDN18.2 was heterogeneously expressed in gastric cancer tissues. (A) Histogram of accumulative IHC score of CLDN18.2 and NKG2DLs in microarray of gastric cancer. (B) Frequency of CLDN18.2-positive, NKG2DL-positive, and dual positive tissues in microarray of gastric cancer. (C) Representative images of gastric carcinoma tissues, stained by CLDN18.2 (top row), ULBP1 (middle row) and ULBP3 (bottom row).(D) Tumor volume of gastric carcinoma in PDX model was surveyed at different time points after CAR-T infusion. (E). Tumor tissues in vehicle and CLDN18.2 CAR-T groups were stained by indicated antibodies at the endpoint of experiment, scale bars=500μm. Bars show mean ± SEM. *, P < 0.05; **, P <0.01; ***, P < 0.001.

To validate the consequence of the heterogeneity of tumor antigen expression in the context of CAR-T therapy, we developed a second-generation CAR-T against CLDN18.2. Our CLDN18.2 CAR-T could specifically and highly effectively kill CLDN18.2 positive cells in vitro (Supplementary Fig. 1A-F). Moreover, the CLDN18.2 CAR-T could eliminate the tumors in a CLDN18.2-highly expressed NUGC4-Luc xenograft model in immunocompromised mice. In contrast, CLDN18.2 CAR-T was much less efficacious in a gastric PDX model with heterogeneous expression of CLDN18.2 (Fig.1D). Analysis of the PDX tumors by IHC staining of CLDN18.2 and NKG2DLs showed that CLDN18.2 expression in the tumor treatment by CLDN18.2 CAR-T was largely negative or low expressed, suggesting the resistance of CLDN18.2-targeting CAR-T treatment due to the loss of the targeting antigen. Importantly, NKG2DLs were still homogenously expressed in the tumor tissues of the CAR-T treatment. All these results demonstrated that the escape of CLDN18.2 expression by tumor cells is one of the causes that influences the anti-tumor efficacy of conventional CLDN18.2-targeting CAR-T cells, and that co-targeting NKG2DLs might be one of the solutions to tackle this problem.

### SNR enhances CLDN18.2 CAR-T cytotoxicity and multiple cytokine secretion in vitro

The results presented herein demonstrate that dual targeting of CLDN18.2 and NKG2DL might greatly enhances the recognition range of CAR-T cells in gastric cancer. To achieve this, we developed synthetic NKG2D receptors (SNRs) by fusing the intracellular domains of DAP10 and DAP12 to the extracellular domain of NKG2D, which were linked by CD8 hinge and transmembrane domains. The SNR was then coupled to the second-generation CLDN18.2 CAR via a 2A self-cleaving peptide (Fig.2A), resulting in efficient transduction of T cells at a rate of 95%, compared to 38% for conventional CLDN18.2 CAR (Fig.2B). Then, we utilized a CLDN18.2 expressed gastric cancer cell line NUGC4 and assessed the activities of SNR CAR-T by coculturing them. Both conventional CAR-T and SNR CAR-T efficiently lysed the CLDN18.2 positive target cells (Fig.2C). To further test the dual-targeting activity of our SNR CAR-T targeting NKG2DLs, we utilized CLDN18.2-negative RKO and A431 cell lines, which express high levels of NKG2DLs for target cells. We transduced these cell lines to generate double-positive cells and mixed them with their parental lines at a 1:1 ratio (Supplementary Fig.2A and 2B). We found that only the SNR CAR-T cells could kill both parental and CLDN18.2-overexpressing A431 and RKO cells (Fig.2D-E). To evaluate CAR-T cytokine secretion, Raji-MICA and Raji-CLDN18.2 were used as target cells to stimulate CAR-T. Raji cells were negative for both CLDN18.2 and NKG2DLs. We found that conventional CAR-T did not release any cytokines, whereas SNR CAR-T exhibited stronger cytotoxicity against Raji-MICA, and secreted higher levels of multiple cytokines, including IL-2, TNFα and IFN-γ (Fig.2F). These results suggested that SNR CAR-T had the dual-targeting activity to kill both antigen-single and double positive cancer cells, which highlights their capability to overcome the heterogeneity of tumor cells.

**Fig.2.**
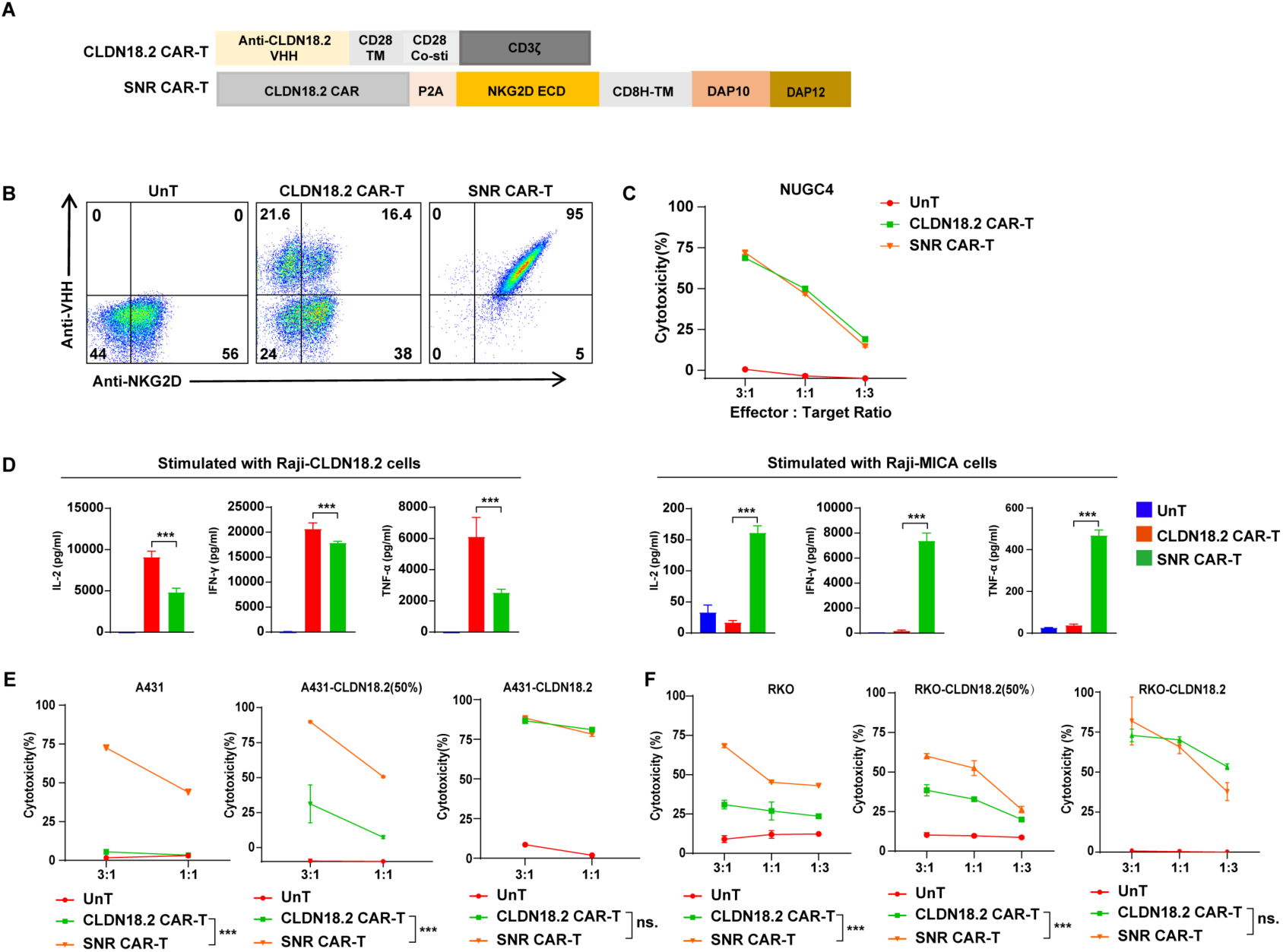
CLDN18.2 CAR-SNR-T could target multiple cancer cells in vitro. (A) Schematic construction of CLDN18.2 CAR-T and SNR CAR-T. (B) Expression of CAR and NNKG2D in different CAR-T were analyzed by flow cytometry. (C) Cytotoxicity of CLDN18.2 CAR-T and SNR CAR-T against gastric cancer cell line NUGC4. (D) Multiple cytokines secretion was analyzed after CAR-T incubation with Raji-CLDN18.2 or Raji-MICA. (E) Cytotoxicity of CLDN18.2 CAR-T and SNR CAR-T against multiple tumor cell lines including A431, A431 and A431-18.2 mixed cells, and A431-18.2. (F) Cytotoxicity of CLDN18.2 CAR-T and SNR CAR-T against multiple tumor cell lines including RKO, RKO and RKO-18.2 mixed cells, and RKO-18.2. Error bars represent mean + SEM. *, P < 0.05; **, P <0.01; ***, P < 0.001.

### SNR enhance the memory phenotype and suppress exhaustion marker expression of CAR-T

We investigated the impact of SNR on the cellular phenotypes of CLDN18.2 CAR-T. To reveal gene expression between SNR CAR-T and conventional CAR-T, we found that SNR CAR-T were transcriptionally distinct from conventional CAR-T from RNA sequencing, with more than 1000 genes differentially expressed in the resting condition (Fig.3A). Among these differentially expressed genes, exhaustion related genes (*EOMES, CD160, LAG3, CTLA4, NFATC4, TOX2*) and activation related genes (*TNFRSF9, TNFSF9, IL2RA, CD69, CD38, TNFRSF4*, *TNFSF4*) were significantly down-regulated in SNR CAR-T cells. Moreover, SNR CAR-T showed higher expression of a subset of T cell memory related genes including (*TCF7, SELL, CD27, CNR2, PDE9A, CTSC, PECAM1, LEF1* (Fig.3A). Further, Gene set enrichment analysis (GSEA) also confirmed that SNR could reduce T cell exhaustion and activation, while enhancing memory formation of CAR-T in unstimulated situation. (Fig.3A) ^26^. To validate the findings from transcriptomic analysis, we measured the cell surface expression of T cell memory and exhaustion markers by flow cytometry (FCM). Consistent with results from gene sequencing, we found that the proportion of Tscm subsets and CD27 expression were significantly higher, and the percentage of PD-1-positive cells were significantly lower in CLDN18.2 CAR-T co-expressing SNR (Fig.3B and 3C). To further characterize the phenotypes of CAR-T at single cell level, we used the CyTOF (cytometry by time of flight) to analyze the expression profile of our T cells in rest condition (Fig.3E). SNR CAR-T showed increased expression of CD62L and CD45RA (Fig.3F and 3G), consistent with the results by FCM (Fig.3B). In term of the T cell activation and exhaustion, SNR CAR-T expressed reduced levels of CD25, CD38, CD39, and PD1 (Fig.3G). Interestingly, a subset of CD8 positive cells with high expression of CD39, CD56 was noted in the conventional CAR-T cells, which were absent in SNR CAR-T (Fig.3G). CD8 T cells with CD39 and CD56 high expression might be dysfunction or terminal exhausted and have inhibitory capacity^27^. A subset of CD4 positive cells with high expression of CD25 and PD1 was noted in conventional CAR-T cells, which was absent in SNR CAR-T. Taken together, these results suggest that SNR CAR-T have the higher memory phenotype cells and reduced exhaustion.

**Fig.3.**
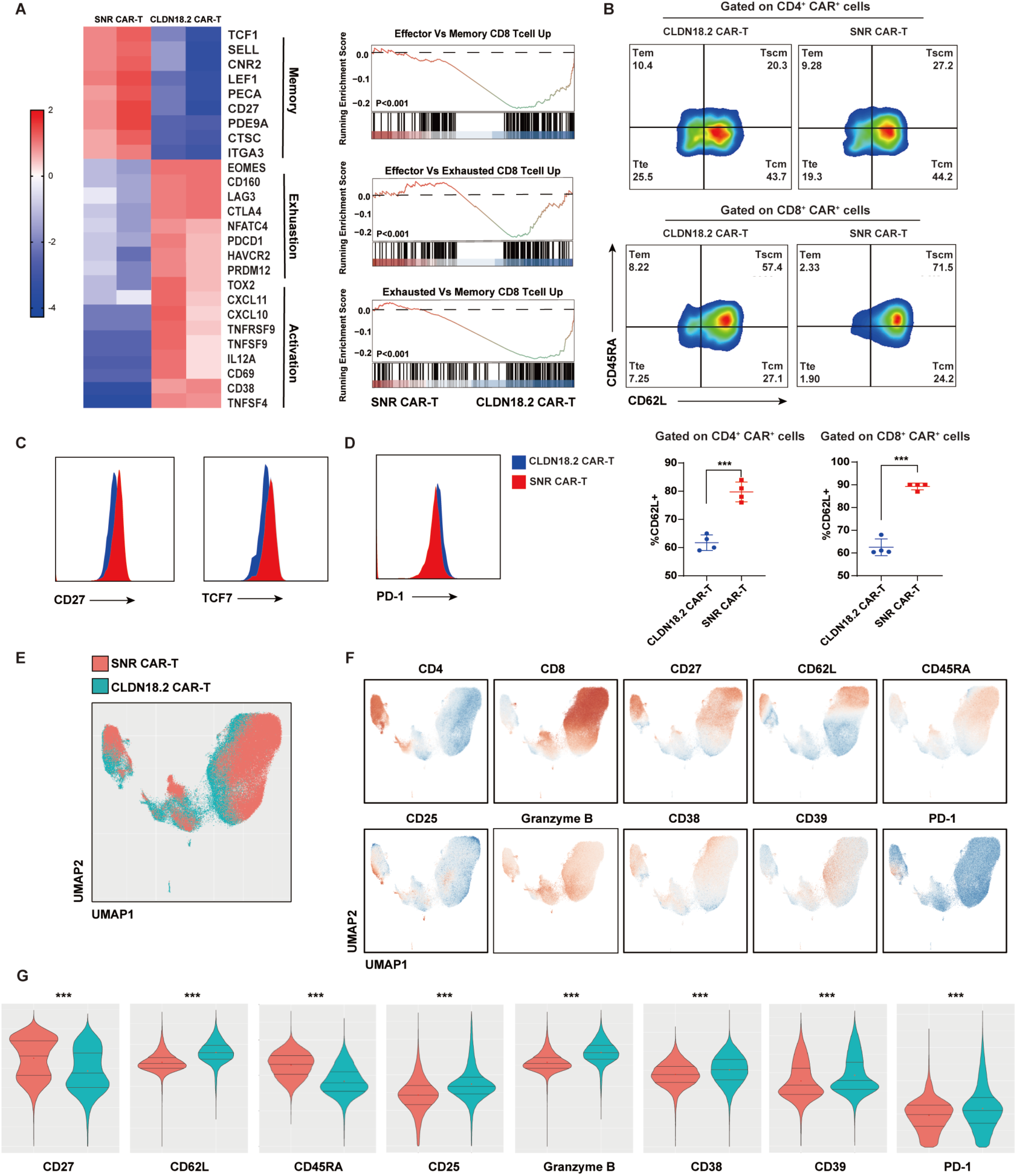
Expression of SNR on CLDN18.2 CAR-T promote CAR-T potency in memory differentiation and reduce exhaustion. (A) Heatmap displayed the differentially expressed genes (DEGs) that are related to T cell activation, memory formation and exhaustion between CLDN18.2 CAR-T and SNR CAR-T. Gene set enrichment analysis (GSEA) of pathways including: Effector VS Memory CD8 T cell up, Effector VS Exhausted CD8 T cell up and Exhausted VS Memory CD8 T cell up between CLDN18.2 CAR-T and SNR CAR-T. (B) Proportion of different subsets of T cells including stem like memory T cells (Tscm), central memory T cells (Tcm), effector memory T cells (Tem) and terminal differentiated T cells (Tte) between CLDN18.2 CAR-T and SNR CAR-T. Percentage of CD62L^+^ T cell between CLDN18.2 CAR-T and SNR CAR-T after stimulation with NUGC4. (C) TCF7 and CD27 expression in CAR-T cells in rest condition was shown. (D) PD-1 expression in CAR-T cells was analyzed in rest condition. (E) UMAP plot of total cell population grouped by CLDN18.2 CAR-T and SNR CAR-T. (F) UMAP plots of CD4, CD8, CD27, CD62L, CD45RA, CD25, Granzyme B, CD38, CD39, and PD1. (G) Violin plot showed relative expression of CD4, CD8, CD27, CD62L, CD45RA, CD25, Granzyme B, CD38, CD39, and PD1 between CLDN18.2 CAR-T and SNR-CAR T. Error bars represent mean + SEM. *, P < 0.05; **, P <0.01; ***, P < 0.001.

### SNR increases anti-tumor efficacy, the expansion and infiltration of CAR-T cells in vivo

To investigate whether the SNR-enhanced functional activities observed in vitro could translate to improved anti-tumor efficacy in vivo, we inoculated two CLDN18.2-high-expressing cancer lines, NUGC4-Luc and MIAPaCa2-CLDN18.2, to generate xenograft tumor models in immunodeficient mice and dosed them with CLDN18.2 CAR-T or SNR CAR-T. Both CLDN18.2 CAR-T or SNR CAR-T can control tumors efficiently and that the size of tumors in SNR CAR-T treated group were significantly smaller or even completely eradicated (Fig.4A and 4C). Both groups of CAR-T were well-tolerated, and there was no evidence of toxicity or significant decrease in the body weight of the mice (Fig.4B and 4D). H&E staining on different organs after CAR-T infusion also indicated that SNR CAR-T didn’t cause tissue damage (Fig.4H). IFN-γ production and CAR-T cell expansion was evaluated at day4, day11 and day18 after the infusion of CAR-T cells. Our data showed that the SNR CAR-T treated mice had significantly higher levels of IFN-γ in their plasma than the conventional CAR-T-treated mice at 2 out of 3 time points (Fig.4E). Moreover, the expansion of SNR CAR-T was more robust than that of control CAR-T at two early time points, reaching its peak at day 11 (Fig.4F). However, no T cell could be found in CLDN18.2 CAR-T-treated mice at day 11 (Fig.4F). Collectively, these findings suggest that SNR enhances the expansion and anti-tumor activity of CAR-T cells in vivo. We further investigate whether SNR could improve tumor infiltration of CAR-T. We used immunohistochemistry (IHC) against human CD45 to detect the distribution of CAR-T in tumors harvested from MIA-Paca2 CLDN18.2 tumor-bearing mice ten days after treatment with CAR-T cells. We found very few T cells were inside the tumor nest and most of T cells were localized in the stroma or the margin surrounding the tumors in the CLDN18.2 CAR-T group (Fig.4G). In contrast, tumors from mice treated with CAR-T cells co-expressing SNR showed intense infiltration of CAR-T cells across all the regions of tumor (Fig.4G). The difference in tumor infiltrating T cells was further illustrated by co-staining CLDN18.2 as tumor marker, human CD4 and CD8 (Fig.4I and 4J). Results suggested that CD4 and CD8 subsets of SNR CAR-T showed stronger proliferating state in tumors than conventional CAR-T (Fig.4I and 4J).

**Fig.4.**
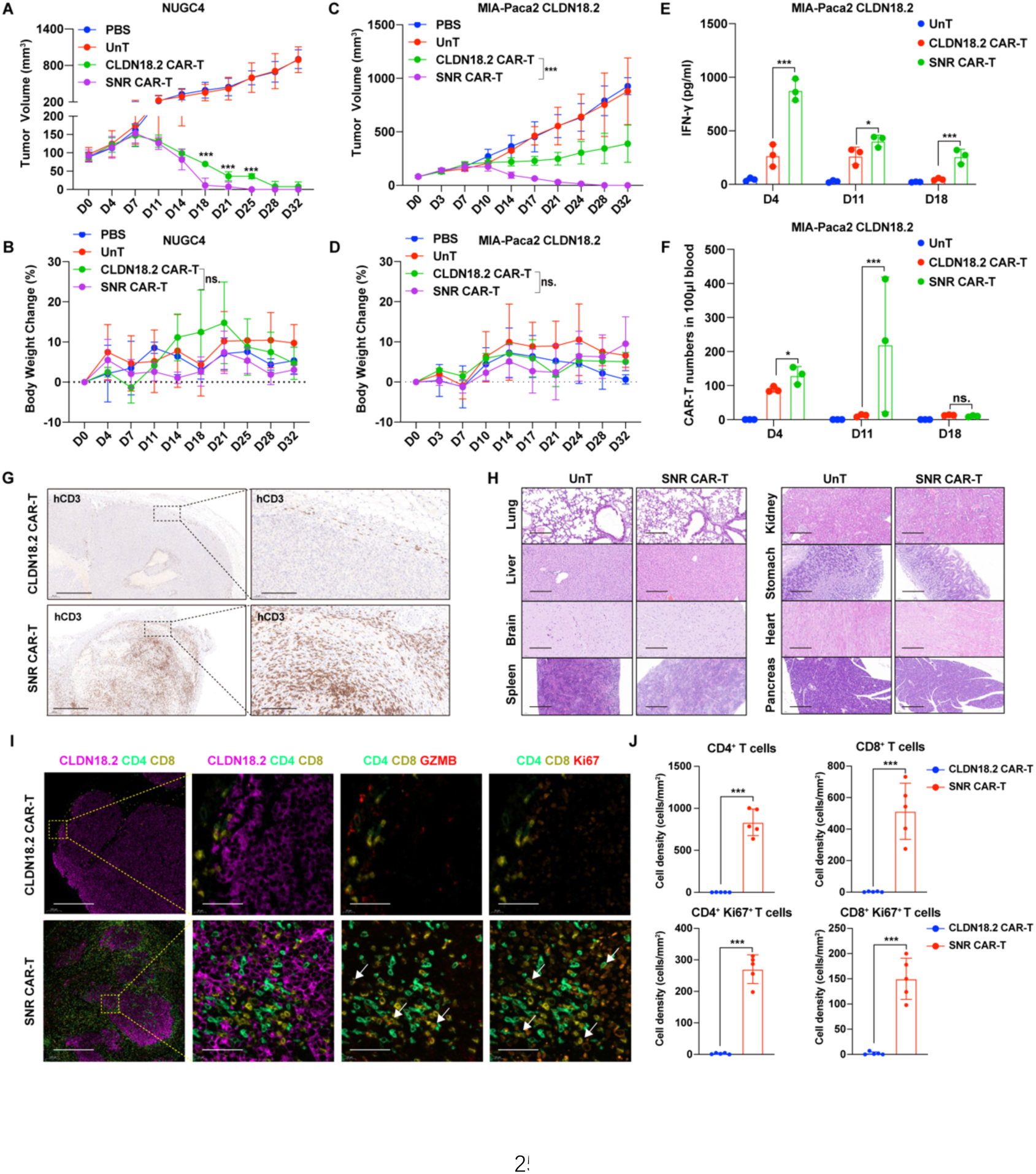
CLDN18.2 CAR-SNR-T cell improved CLDN18.2-dependent antitumor efficacy in vivo. (A) Tumor volume of NUGC4 cell derived xenograft at different time points post CAR-T injection. NUGC4 cells (5.0×10^6^) were subcutaneously implanted into NSG mice. The mice were intravenously infused with 1×10^6^ CAR-T cells (n=3). (B) Average body weight change of mice from three groups within 32 days. (C) Tumor volume of MIA-Paca-2 derived xenograft at different time points after CAR-T injection were measured. MIA-Paca-2 cells (CLDN18.2 positive; 1.0×10^6^) were subcutaneously implanted into NSG mice and (D) the average body weight change was monitored. (E) IFN-γ concentration (pg/ml) of peripheral blood in mice bearing MIA-Paca-2 tumors were detected on day4, day11 and day18. (F) The absolute number of CAR-T in 100μL blood were detected on day4, day11 and day18 after CAR-T infusion. (G) MIA-Paca-2 (CLDN18.2 positive) tumors were engrafted subcutaneously, treated with CAR-T cells, and analyzed by IHC for T cell infiltration (anti-human CD3). CLDN18.2 CAR-T failed to penetrate the tumor and accumulated in tumor edges (top row). SNR CAR-T resulted in substantially increased T cell infiltration into tumor core (bottom row). Scale bars are 500 μm and 50 μm. (H) HE staining of different organs following SNR CAR-T infusion. (I) Multiplex-IHC staining of CD4, CD8, GZMB, Ki67 were performed to investigate the infiltration and status of tumors treated by CLDN18.2 CAR-T or SNR CAR-T. (J) Statistics data of CD4, CD8, CD4^+^ Ki67^+^, CD8^+^ Ki67^+^cell counts in tumor. Error bars represent mean + SEM. *, P < 0.05; **, P <0.01; ***, P < 0.001.

Overall, these findings suggested that SNR significantly enhance the anti-tumor activity of CAR-T cells in xenograft tumor models with homogeneously expressed antigen and dramatically increase the T cell infiltration into the tumors.

### SNR armored CAR-T overcome the tumors with target heterogeneity in vivo

Due to the antigen heterogeneity of target tumor antigens in clinical, we generated a in vivo tumor model by mixing CLDN18.2-negative/NKG2DLs-high parental A431 cells with CLDN18.2-overexpressing A431 cells at a ratio of 7:3. As expected, the SNR CAR-T significantly inhibited tumor growth (Fig.5A) and showed significant T cell expansion at day 9 (Fig.5B). In contrast, conventional CAR-T failed to control tumor growth and were unable to expand following infusion (Fig.5B). Further, we tried to determine whether the enhanced anti-tumor efficacy observed in cell-derived xenograft (CDX) tumor models could be also recapitulated in human-derived tumors in vivo. We compared the activity of SNR CAR-T in patient-derived xenograft model (PDX) with the conventional counterpart. The IHC of the PDX tumor sample for the expression of CLDN18.2 and NKG2DL showed uneven expression of CLDN18.2 with same area of negative staining (Fig.1E) and intense expression of some NKG2DL (Fig.1D). We found that both CAR-T cells were very potent to control the tumor growth very efficiently and that SNR CAR-T demonstrated the trend of better T cell expansion and tumor growth control (Fig.5D), compared with the conventional CAR-T.

**Fig.5.**
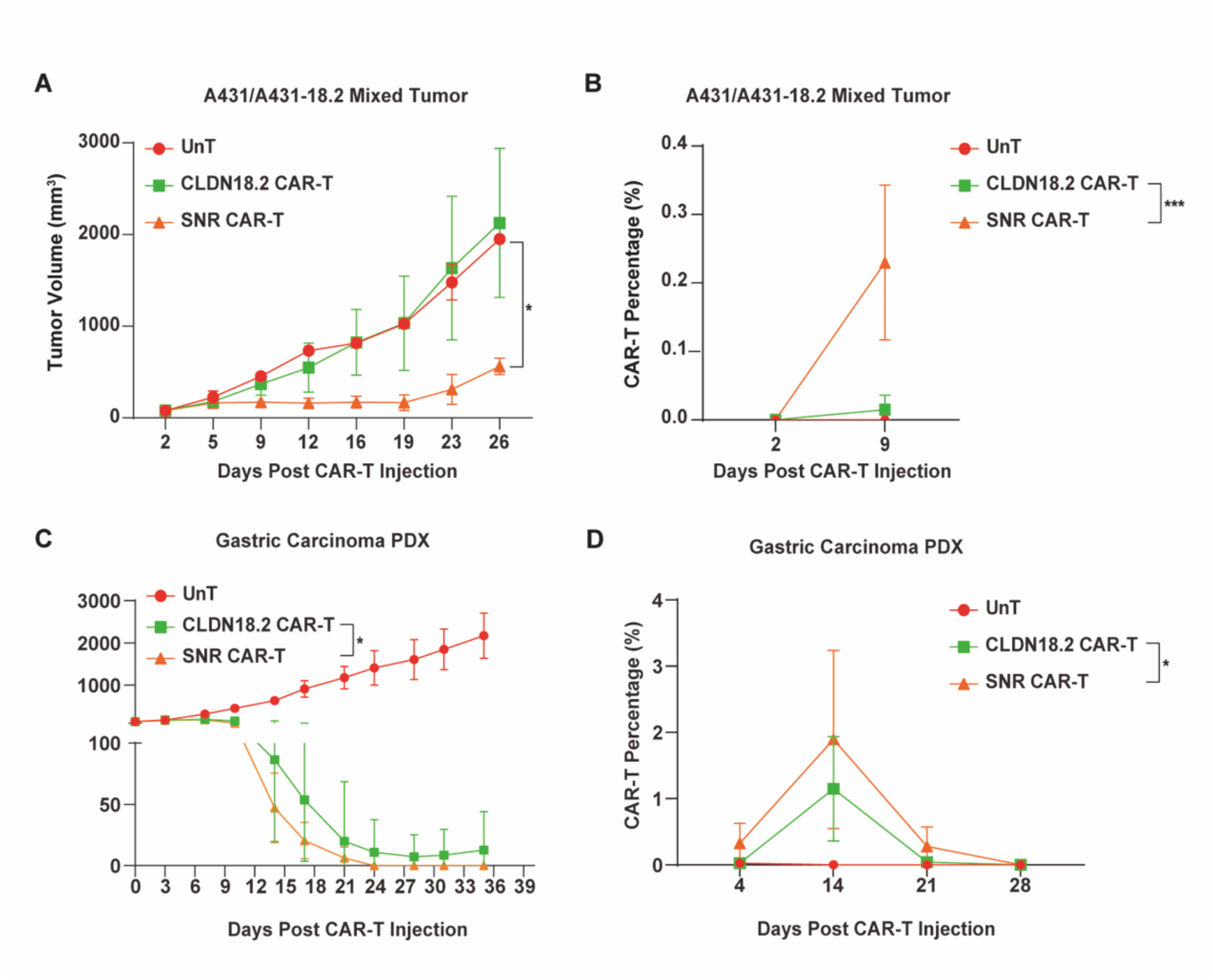
SNR CAR-T had superior antitumor effect with target heterogeneity in vivo. (A) CLDN18.2 heterogenous xenograft model A431/A431-18.2 was constructed and tumor volume was measured. A431 and A431-18.2 was mixed at 7:3 ratio and mixed cells (1.0×10^6^) were subcutaneously implanted into NSG mice. The mice were intravenously infused with 1×10^6^ CAR-T cells. (B) CAR-T percentage in peripheral blood after CAR-T infusion were analyzed by flow cytometry. (C) Tumor volume was measured in mice bearing patient-derived xenograft model (PDX). (D) CAR-T percentage in peripheral blood at determined time points after CAR-T infusion in PDX model. Error bars represent mean + SEM. *, P < 0.05; **, P <0.01; ***, P < 0.001.

Taken together, our results suggest that SNR-armored CLDN18.2 CAR technology confers T cells with the ability to target both tumor-associated antigens and NKG2DLs, which sheds new light on approaches for treating cancer patients with heterogeneous tumor antigen expression.

## Discussion

Despite immunotherapy has shown clinical benefits in advanced gastric cancer (AGC), only a limited number of late phase patients could achieve clinical response^28–31^. Although CAR-T therapy has achieved tremendous progress in hematopoietic malignancies including leukemia, lymphoma, and multiple myeloma, CAR-T therapy targeting solid tumors still faces many obstacles. The heterogeneity of cancer antigen expression is one of the major challenges in the treatment of solid tumors^5,32,33^. Previous study has showed significantly higher infiltration by 20 types of immune cells in the group with low heterogeneity, compared to the group with high heterogeneity scores. And low heterogeneity strength predicted longer overall survival (OS), when compared to those with high scores^34^. Thus, it is necessary to develop a therapeutic strategy to combat the heterogeneity in solid tumors.

It has been shown that NKG2DLs were universally expressed on many solid and hematopoietic malignancies, including gastric cancer^18,35–39^. Consistent with the literatures, we demonstrated that most of gastric cancer tissues were stained positive for at least one NKG2DL by using a commercial gastric cancer tissue array, and that only 38% of samples heterogeneously stained positive for CLDN18.2, in agreement with other publications. Co-targeting both CLDN18.2 and NKG2DLs in the same CAR-T cells could significantly reduce the opportunity of antigen escape of tumor cells.

In this study, we developed a SNR CAR-T to harness the killing activity of NKG2D, a receptor to activate NK cells upon binding to its cognate ligands, such as MICA/B and ULBP1-6, to broaden the therapeutic spectrum of conventional CAR-T. Our SNR is composed of extracellular domain of NKG2D, and intracellular domain of DAP12 and costimulatory domain of DAP10. In addition to kill the CLDN18.2-positive tumor cells, the SNR could guide CLDN18.2 CAR-T cells to lyse NKG2DLs-positive tumor cells and demonstrated synergic effects with CLDN18.2 CAR in vitro and in vivo.

In addition to expand the targeting spectrum of CAR-T, SNR CAR-T also demonstrated the memory and less-exhausted gene expression profiles. Our transcriptomic and proteomic analysis shows Higher expression of memory related genes such as *TCF7, LEF1, SELL* in the SNR CAR-T, indicating that SNR signaling could prevent the differentiation of T cells and tilt the balance toward the memory phenotype. Furthermore, these analysis also found SNR CAR-T expressed less exhaustion genes, such as *LAG3*, *CTLA-4*,and *PDCD1* It has been reported that NKG2D signaling in CD8 T cells is necessary for the development of functional memory cells^40–42^. NKG2D mainly signaling through DAP10 in human CD8 and that DAP10 signaling were demonstrated critical for production of IL-15 and activation of PI3K, which is crucial for survival and homeostasis of memory and memory precursor T cells^43–45^. Thus, the activation of DAP10 signaling might contribute to the enhance of memory formation of CAR-T cells ^25^. Thus, we conclude that the enhancement of memory formation and decrement in exhaustion might be due to the activation of DAP10 signaling pathway through SNR and that SNR CAR-T might have higher proliferation potential and functional activity to kill tumor cells.

We demonstrated SNR can synergize with CAR to eliminate tumors in a NKG2DL-independent manner. SNR CAR-T showed robust T cell expansion in vivo, superior tumor infiltration and anti-tumor efficacy in both models, compared with its conventional counterpart. Part of these enhancement could be due to the reasons that SNR signaling could increase T cell proliferation and decrease their exhaustion as we observed in vitro. However, the precise mechanisms require further investigations. Dual or multiple targeting is currently one of the approaches to tackle the problem of therapeutic resistance developed by cancer cells through antigen escaping. Numerous studies have shown different technologies of dual specific CAR-T systems by expressing 2 CARs in the T cells and demonstrated that CAR-T cells with the capability to target two tumor antigens could significantly improve their anti-tumor activity and decrease the opportunity of antigen-free resistance. These technologies usually target 2 different tumor antigens and need 2 specific antibodies, which significantly increases the difficulty and complexity of the CAR-T development. Another solution to circumvent these hurdles is to take advantage of some receptors expressed by immune effector cells, such as NKG2D, to develop a universal co-targeting CAR. We provided the evidence that SNR armored CLDN18.2 CAR-T has the designed capability to overcome the hurdles of the heterogeneity so that both double-positive and CLDN18.2/NKG2DL single-positive cancer cells can be eradicated in vivo.

In summary, we have developed SNR armored CLDN18.2 CAR-T system to target both CLDN18.2 and NKG2DLs for the treatment of solid tumors. We demonstrated that SNR with DAP10 and DAP12 co-stimulatory domains could improve the memory phenotypes of T cells and increase in vitro and in vivo efficacies against cancer cells or tumors with homogeneous or heterogeneous expression of cancer antigen. Our SNR might have the potential to be a universal platform to arm other CAR-T cells targeting different tumor antigens. Further characterization and clinical development are under the way.

## Data Availability Statement

The authors confirm that the data supporting the findings of this study are available within the article and its supplementary materials.

## Acknowledgments

We thank Yuhan Wang, Dr. Shijia Wang, Dr. Xinzhu Wang from Polaris Biology, Shanghai for performance of mass cytometry experiment and analysis data. We thank Shanghai Model Organisms for raising tumor-bearing mice and performing CAR-T infusion. We thank professor Di Zhu from School of Pharmacy, Fudan University for project design and support.

## Author Contributions

YC, MS and RH designed in vitro and in vivo experiments; YW, YL and HW performed in vitro and in vivo experiments; BZ, MS,YW, RH, HW and SZ collected and analyzed data; YZ constructed plasmids and lentiviral vector; MS, RH, YW and SZ wrote the manuscript.

## Declaration of Interests

M.S., H.W., Y.W., R.H., Y.L., Y.Z. and S.Z. are employees of Suzhou Immunofoco Biotechnology Co., Ltd., for which potential product is studied in this work. The other authors have no competing interests.

---

**Supplementary Fig.1.**
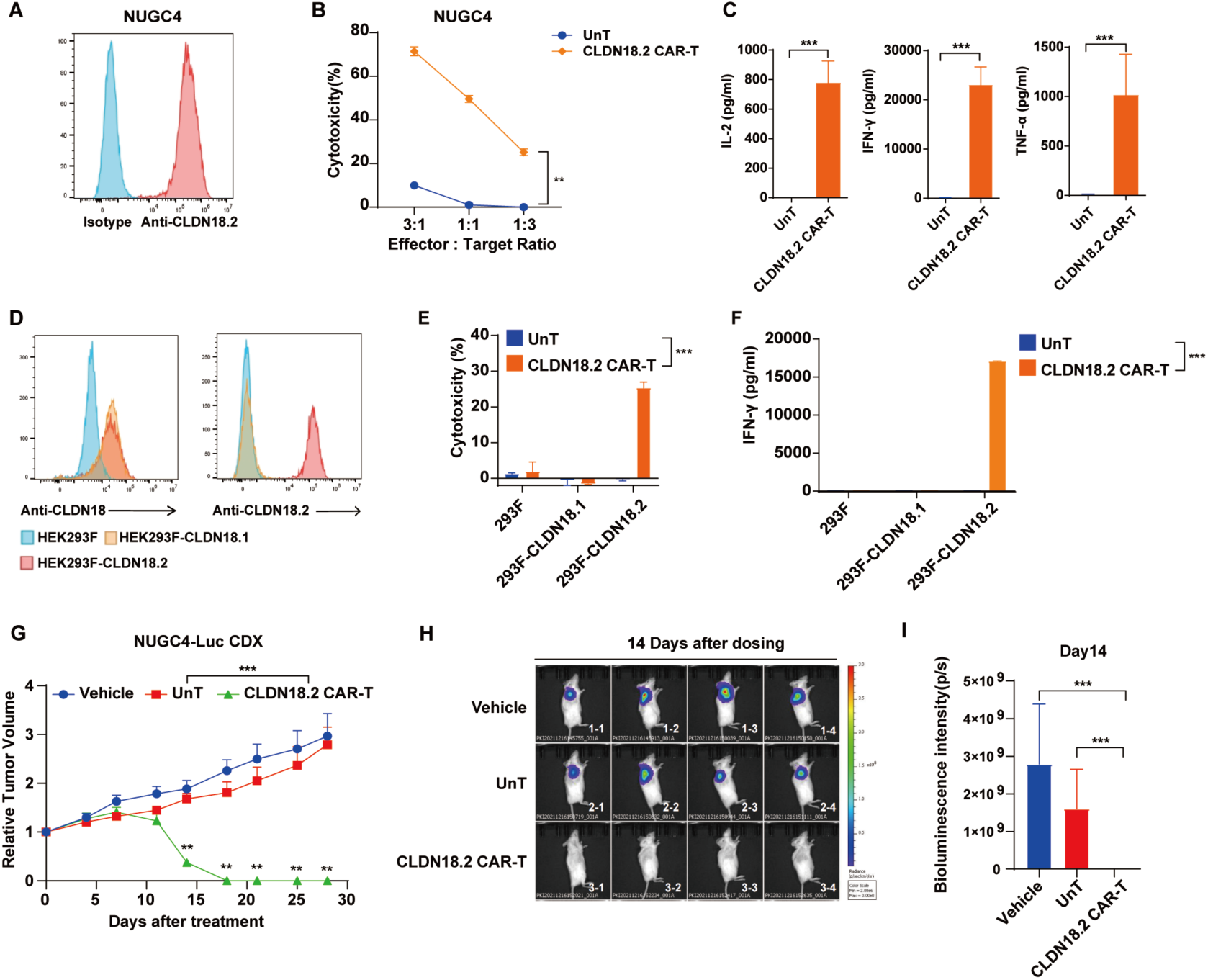
Validation of specificity and efficiency of scFv from CLDN18.2 CAR-T both in vitro and in vivo. (A) CLDN18.2 on NUGC4 cells was identified. (B and C) Killing efficiency of CLDN18.2 CAR-T against NUGC4 and cytokine secretion was detected. (D-F) HEK293F cells that overexpressing CLDN18.1 and CLDN18.2 were contructed, the specificity of CLDN18.2 CAR-T was measured by killing assay and IFN-γ release. (G-I) Antitumor effect of CLDN18.2 CAR-T was evaluated in NUGC4 cell derived-xneograft model, tumor volume and tumor-related bioluminescence intensity were quantified.

**Supplementary Fig.2.**
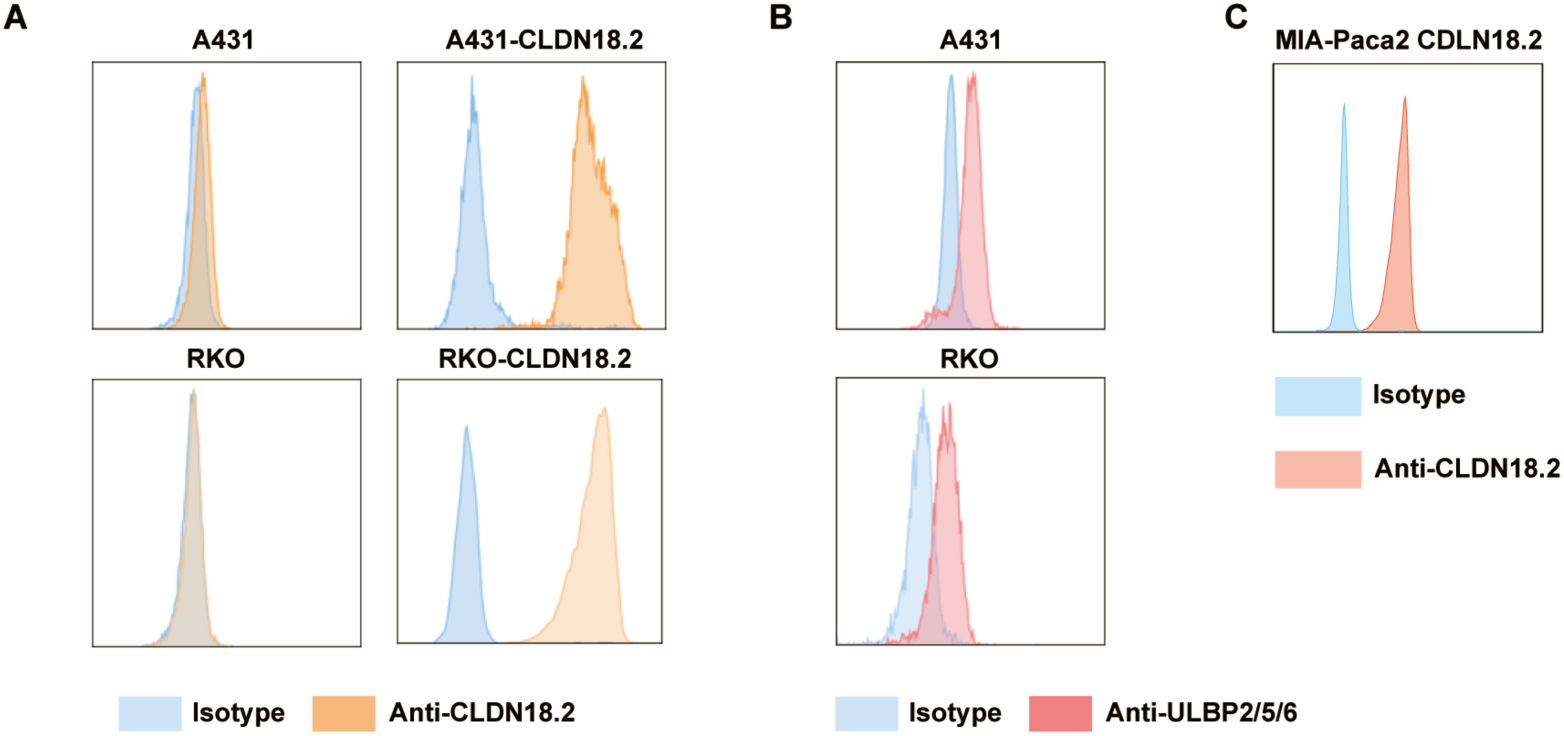
Construction of different CLDN18.2 overexpressed cell lines. (A) Construction of A431 CLDN18.2 and RKO CLDN18.2 cell line. CLDN18.2 expression in each cell lines were detected via flow cytometry. (B) NKG2DLs expression in A431 and RKO. (C) Construction of MIA-Paca2 CLDN18.2 cell line.

